# The emergence of tuning to global shape properties of radial frequency patterns in the ventral visual pathway

**DOI:** 10.1101/2023.03.01.530430

**Authors:** Samuel J. D. Lawrence, Elisa Zamboni, Richard J. W. Vernon, André D. Gouws, Alex R. Wade, Antony B. Morland

**Affiliations:** Department of Psychology, University of York, YO10 5DD; York Neuroimaging Centre, University of York, YO10 5NY; York Biomedical Research Institute, University of York

**Author notes:** **Corresponding Author** Professor Antony B. Morland, Department of Psychology, University of York, Heslington, York, YO10 5DD.

**Keywords:** Lateral Occipital Complex, Shape processing, Curvature, MVPA, Representational Similarity Analysis, Retinotopy

## Abstract

Radial frequency patterns - created by sinusoidal modulations of a circle’s radius - are processed globally when radial frequency is low. These closed shapes therefore offer a useful way to interrogate the human visual system for global processing of curvature. Radial frequency patterns elicit greater responses than those to radial gratings in V4 and more anterior face selective regions of the ventral visual pathway. This is largely consistent with work on non-human primates showing curvature processing emerges in V4, but is evident also higher up the ventral visual stream. Rather than contrasting radial frequency patterns with other stimuli, we presented them at varied frequencies in a regimen that allowed tunings to radial frequency to be derived from 8 human participants (3 female). We found tuning to low radial frequency in lateral occipital areas and to some extent in V4. In a control experiment we added a high frequency ripple to the stimuli disrupting the local contour. Low frequency tuning to these stimuli remained in the ventral visual stream underscoring its role in global processing of shape curvature. We then used representational similarity analysis to show that in lateral occipital areas the neural representation was related to stimulus similarity, when it was computed with a model that captured how stimuli are perceived. We show therefore that global processing of shape curvature emerges in the ventral visual stream as early as V4, but is found more strongly in lateral occipital regions, which exhibit responses and representations that relate well to perception.

## Introduction

Processing of increasingly complex spatial features occurs up the hierarchy of the ventral visual pathway. At the start of the pathway, V1 processes orientation (Hubel & Wiesel, 1968), while areas high up the pathway respond selectively to real-world objects (Tanaka, 1996), (Grill-Spector et al., 2001) or faces (Desimone et al., 1984; Kanwisher et al., 1996; Tsao et al., 2006). The intermediate levels of the pathway process spatial features such as curvature (Kourtzi & Connor, 2011), which has long been seen as part of pattern vision (Riggs, 1973) and is the focus of the present paper.

Pioneering work assessed responses in the intermediate area V4, where many cells had response preferences to polar and hyperbolic stimuli, which exhibited curvature, over the straight lines of cartesian stimuli (Gallant et al., 1993, 1996). Others have shown in experiments using angles and curves (Pasupathy & Connor, 1999) and simple curved shapes (Pasupathy & Connor, 2001) how V4 neurons are selective to boundary conformations (Pasupathy et al., 2020). V4 neurons also respond to other information (Zeki, 1973) with domains specific to curvature, orientation and colour being evident (Conway et al., 2007; Hu et al., 2020; Roe et al., 2012; Tang et al., 2020). Experiments using naturalistic visual stimuli, from which curvature was quantified, have shown that curvature processing is not limited to V4 and that two regions more anterior in the ventral visual pathway also respond to curvature in macaque (Yue et al., 2014). V4 has therefore been referred to as ‘curvature emergent’ as areas antecedent to it do not appear to process curvature (Hu et al., 2020).

In human a role for V4 and other extrastriate areas in processing curvature has also been found. Wilkinson et al found greater responses in V4 to concentric than parallel patterns, suggesting that there was a preference for curvature in V4 (Wilson et al., 1997). However, it was only in the more anterior fusiform face area (FFA) that responses to the concentric, curved patterns were greater than the straight lined radial patterns. A preference for concentric curvature (defined in arrays of Gabors) was found in V4, but also earlier in V3 (Dumoulin & Hess, 2007). We have found that shape curvature representations emerge in areas LO1 and LO2, which could be considered as one step further up the object processing pathway than V4 (Vernon et al., 2016). Yue et al (Yue et al., 2020) found preferences to curvature in V4 and also to some extent in V3 and also in regions that were more anterior in the ventral visual pathway, where more complex aspects of curvature were processed (Yue et al., 2014). The Lateral Occipital Complex (LOC) (Grill-Spector et al., 2001; Malach et al., 1995) has also been found to have a shape representation based upon shape features (Drucker & Aguirre, 2009; Haushofer et al., 2008; Op de Beeck, Haushofer, et al., 2008). There is, therefore, reasonably broad agreement between there being an emergence of curvature processing in the human and macaque brain in V4 and that curvature information is processed further in regions higher up the ventral pathway.

The stimuli used to investigate curvature have been justifiably varied with configurations spanning highly controlled, narrow band stimuli (Dumoulin & Hess, 2007) to more naturally realistic images (Yue et al., 2020). Here, we use radial frequency patterns, which have been presented in studies of processing of shape curvature in humans (Wilkinson et al., 2000) and non-human primates (Tang et al., 2020). Radial frequency patterns have also been a mainstay of psychophysical literature that has shown exquisite sensitivity to (Wilkinson et al., 1998) and global processing of low radial frequency stimuli (Bell et al., 2007b; Hess et al., 1999; Jeffrey et al., 2002; Lawrence et al., 2016; Loffler et al., 2003). We predicted therefore that response preferences to, and representations of, low radial frequencies will emerge in the ventral visual pathway.

## Methods

### Rationale

The aim of the study was to understand the tuning to radial frequency in the following regions of interest; V1-V4, LO1, LO2 and LO. While these regions together do not correspond to the complete ventral visual pathway, we predicted that global processing of shape would emerge in at least one of these regions. We followed a paradigm developed by Harvey et al. (Harvey et al., 2013) to allow tuning to a visual parameter, in their case numerosity and in our case radial frequency, to be extracted from brain responses to a specific and effective stimulus regimen. The approach lends itself to in-depth assessment of relatively few individual participants - in our case eight, like the study by Harvey et al (2013) - by acquiring a large amount of data (~5 hours) from each individual. The paradigm is also well suited to establishing whether there are topographic mappings of the parameter under investigation. In our study, however, we found no strong evidence of a topographic mapping of radial frequency. Our investigation therefore followed the general approach of evaluating univariate and multivariate responses within the regions of interest that were identified in individuals from retinotopic mapping and functional localizer experiments to establish what radial frequency tuning properties those regions exhibited. In the sections that follow, the participants we tested are described followed by the MRI details, and then the procedures for retinotopic mapping, LOC localizer and RF tuning experiments are given.

### Participants

Eight participants (Mean age 28.00, SD 4.75; 5 Males) were recruited from the University of York Psychology department. All participants had normal or corrected-to-normal visual acuity, were naive to the aim of the study, and gave written informed consent. All participants completed 5.25 hours of scanning in total, including a structural session, a retinotopic mapping session, an LOC localiser session, and two main radial frequency tuning sessions. The study was approved by the York Neuroimaging Centre Ethics Committee in accordance with the Declaration of Helsinki.

### MRI

#### fMRI data acquisition

All imaging data were acquired on a GE 3-Tesla Sigma HD Excite scanner using a 16-channel half-head coil to improve signal-to-noise in the occipital lobe. Acquisition parameters and analysis procedures for each session are described below. Across all experiments stimuli presentation were controlled using Matlab and Psychophysics toolbox (Brainard, 1997). Stimuli were presented using a projector and mirror setup (Dukane Image Pro 8942 LCD projector, pixel resolution 1280×1024, 60 Hz frame rate) at a viewing distance of 57 cm.

#### Structural scans

We acquired three, 16-channel, T1-weighted anatomical images (TR = 7.8ms, TE = 3.0ms, TI = 600ms, voxel size = 1×1×1mm^3^, flip angle = 12°, matrix size 256×256×176, FOV = 25.6cm), one 8-channel T1-weighted anatomical image to aid alignments (TR = 7.8ms, TE = 2.9ms, TI = 450ms, voxel size = 1.13×1.13×1mm^3^, flip angle = 20°, matrix size 256×256×176, FOV = 29cm) and one T2*-weighted fast gradient recalled echo scan (TR = 400ms, TE = 4.3ms, voxel size = 1×1×2mm^3^, flip angle = 25°, matrix size 128×128, FOV = 26cm).

T1-weighted anatomical data was used for coregistration and gray-white matter segmentation. To this end, the three 16-channel T1 scans were aligned and averaged together. We then divided this average by the T2*-weighted data to improve grey-white matter contrast and partially correct for the signal drop-off caused by use of a half-head coil. The resulting average T1 was then automatically segmented into grey and white matter using FreeSurfer (Fischl, 2012) and, where necessary, manually improved after visual inspection using ITKGray (https://web.stanford.edu/group/vista/cgi-bin/wiki/index.php/ItkGray).

### Retinotopic mapping

Each participant completed 6 wedge scans (size: 90°, rotating counterclockwise) and 2 expanding ring scans for this session (TR = 3000ms, TE = 30ms, voxel size = 2×2×2mm^3^, flip angle = 90°, matrix size 96×96×39, FOV = 19.2cm). Each scan contained 8 cycles of wedges/rings, with 36s per cycle, traversing a circular region of radius 14.34deg. Both wedges and rings were high contrast (>98%, 400cdm^−2^) checkerboard stimuli that flickered at a rate of 6Hz. Participants were instructed to attend to a central red fixation cross throughout the scans.

We performed standard analysis on the retinotopy data (Wandell et al., 2007), as specified previously (Baseler et al., 2011). For each participant, we identified V1-V3 as our early retinotopic regions, V4 for comparison with Macaque literature, and LO1/LO2 (Larsson & Heeger, 2006) as potential transitionary regions between retinotopic and object-based representations (Silson et al., 2013; Vernon et al., 2016). All ROIs were identified in both hemispheres for all participants, however as we had no *a priori* reason to suspect hemispheric differences, all ROIs were collapsed across hemispheres.

### LOC localiser

To identify the Lateral Occipital Complex (LOC), each participant performed three 8min localiser scans (imaging parameters were identical to those used for retinotopy). Each scan comprised sixteen interleaved objects and scrambled objects blocks in an ABAB design format. Each block lasted 15s, with one stimulus presented per second (0.8s presentation, 0.2s interstimulus interval). To ensure participants attended to the stimuli throughout the session, they performed a one back task in which there could be one, two or no repeated items within a given block, while maintaining fixation at a red central cross. All stimuli were presented on a full screen mid-grey background (200cdm^− 2^) and there were no baseline/rest periods between blocks.

Stimuli comprised 225 easily recognisable greyscale object images. Background information was removed and image histogram was equalised. All objects were set to subtend 4×4 degrees visual angle on average. The scrambled object images were obtained by splitting the object images into squares of 0.8×0.8 degrees of visual angle in size. Any squares lying within the convex hull of the object were then randomly permuted and rotated. This removed any coherent form, while preserving both the coarse global shape profile and local details. Furthermore, a Gaussian filter (SD 1px) was applied to both object and scrambled image sets.

Localiser data were analysed using FEAT (FMRI Expert Analysis Tool; Worsley, 2001). At the first (individual) level we removed the first three volumes and used a high-pass filter cutoff point of 60s to correct for low-frequency drift. Spatial smoothing was set to 4mm and FILM prewhitening was used. Head movements were corrected for (MCFLIRT) and the resultant six motion parameters were entered as confound covariates in the GLM model. To combine data within participant we ran fixed-effects analysis with cluster correction (Z > 5.0, *p* < .001).

Grey matter restricted, cluster-corrected significant activity from the LOC localiser was rendered on the individual surface of each participant. LO was manually defined as the largest cluster in each hemisphere, avoiding overlap with the retinotopically identified nearby LO2 region. These clusters were then mapped back to grey matter, collapsed across hemispheres.

### Stimulus localiser

To restrict ROIs to the stimulus representation, participants performed a localiser scan (imaging parameters were identical to those used in the retinotopic mapping session). Each scan started with a 10s fixation period, followed by 5 blocks of standard and 5 blocks of control radial frequency stimuli (see *Radial Frequency Stimuli*), presented pseudo-randomly and interleaved with fixation blocks. In each block, the seven individual stimuli (RF2-7 and RF10 patterns) were presented twice, resulting in a 14s block (stimuli were on screen for 0.8s, followed by a 0.2s interstimulus interval). Patterns were presented centrally against a mid-grey background, and participants were instructed to fixate on a black central cross while performing an oddball task on the RF stimuli (1 in 10 were contrast reversed).

Data analysis followed the description in the LOC localiser section, with the exception that 5 dummy volumes were removed to allow the scanner to reach a steady state and the high-pass filter cutoff point was set to 84s. We used cluster correction (Z > 3.1, *p* <.05) to identify significant voxels with which to restrict ROIs.

This localiser primarily affected V1-V3, keeping an average of 28.6%, 42.2% and 52.2% of voxels respectively. The V4 (89.1%), LO1 (96.3%), LO2 (85.1%) and LO (88.6%) ROIs were largely preserved.

### Radial Frequency tuning

Each participant completed 2 sessions, each comprising eight, 294s scans. In each session we presented a stimulus set that comprised either a range of radial frequency patterns, we term the ‘RF’ stimulus set, or those same radial frequency patterns with the addition of a high (rf = 20) ripple added to them, a set we term the ‘RF-ripple’ stimulus set. The order of the sessions was counterbalanced across participants. Imaging acquisition parameters were identical to those from the retinotopic mapping session apart from the TR now being set to 2 rather than 3s.

#### RF stimuli

Radial Frequency (RF) patterns (Wilkinson et al., 1998) are defined using the formula:

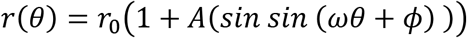

Theta (θ) represents the angles around a circle’s perimeter, allowing the sinusoidal modulation of that perimeter by altering frequency (ω) and amplitude (A; set to 0.1), rotation can be set by altering phase (ϕ). The mean radius (r_0_), governing the average size of the stimulus, was set to 2.5° visual angle.

For the RF stimulus set, we used frequencies 2 to 7, 10 and 20. We also introduced an additional sinusoidal modulation set to a radial frequency of 20 to patterns RF2-7 and RF10 (and RF20 was unchanged) to generate an alternative stimulus set - ‘RF-ripple’. The RF-ripple stimuli allowed the global shape of low RF patterns to be largely preserved but altered the contrast energy of these stimuli, so they were better matched to those with higher radial frequency (see stimulus properties below).

To display the shapes, the contours were rendered against a mid-grey background using the fourth derivative of a Gaussian (Wilkinson et al., 1998) at 50% contrast, yielding a peak spatial frequency of 2 cycles per degree.

#### Stimulus properties

Previous work has shown that spatial frequency and contrast energy content of RF stimuli can vary with both radial frequency and the amplitude. Moreover, variations in these stimulus properties captured a relatively large amount of the multivariate response in visual cortex (Salmela et al., 2016). In the present study, therefore, we attempted to account for and change the relationship between radial frequency and contrast energy and largely equate spatial frequency content of our stimuli.

Figure 2A shows the stimuli we presented; RF stimuli and RF-ripple stimuli in the top and bottom rows respectively. The contrast energy of RF stimuli increases monotonically with radial frequency, but for the RF-ripple stimuli it varies less and is no longer monotonic (Fig. 2B). Because many neurons respond to contrast and our stimuli exhibit variations in contrast a consideration of how a contrast tuned response may register in terms of RF tuning is needed. For the RF stimulus set a contrast tuned response would register tuning to high radial frequency stimuli, while it would be tuned to a lower radial frequency (circa 10) for the RF-ripple stimuli. It is important to note that these tunings are to radial frequencies that are greater than those that are processed globally (<7) and therefore tunings to shape curvature that are behaviorally relevant can be disambiguated from tunings that are driven by sensitivity to contrast.

**Figure 1.**
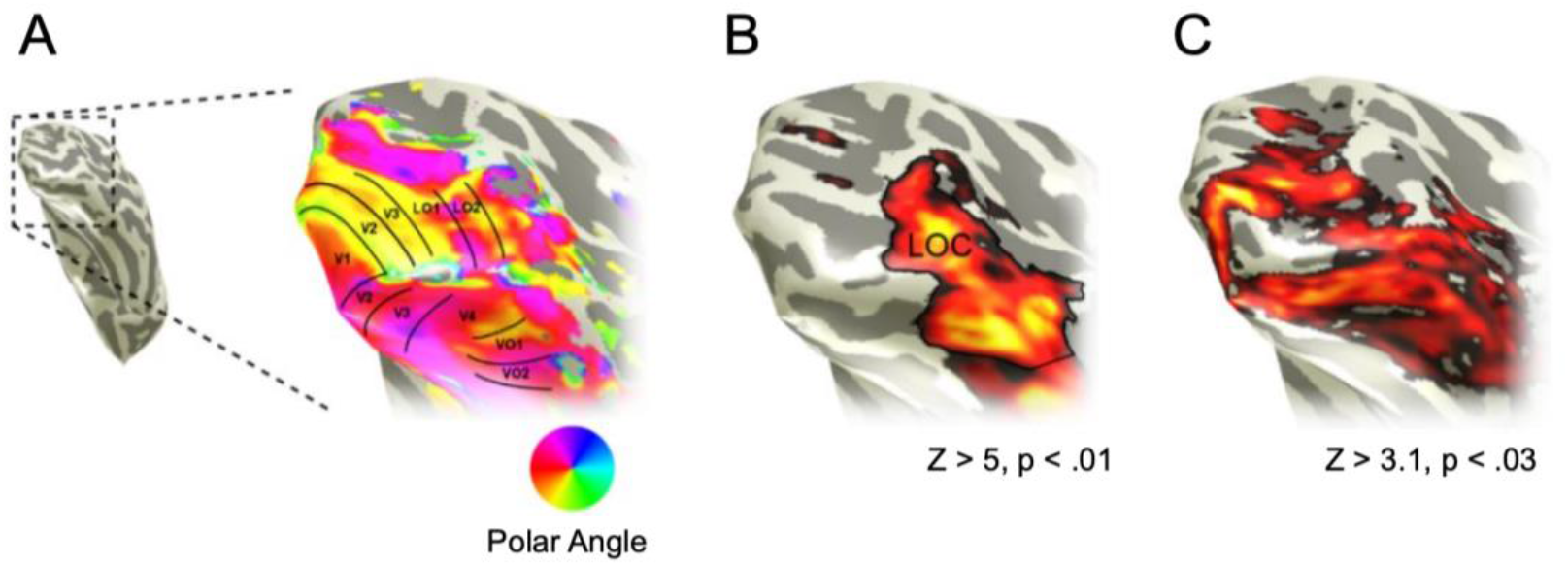
Typical data for an individual participant from the retinotopic mapping, LOC localiser and stimulus localiser experiments for a single subject. (A) Retinotopic mapping was used to identify ROIs V1, V2, V3, LO1, LO2, V4, VO1 and VO2. (B) An objects vs scrambled objects localiser was used to identify LOC as a cluster of object-selective voxels in lateral occipital cortex. We analyzed only the posterior aspect of LOC, which we refer to as LO. (C) A stimulus localiser experiment was used to identify voxels in visual cortex that responded to the stimuli. All ROIs were constrained to only include these voxels to ensure that all analyses only considered voxels within the stimulus representation.

**Figure 2.**
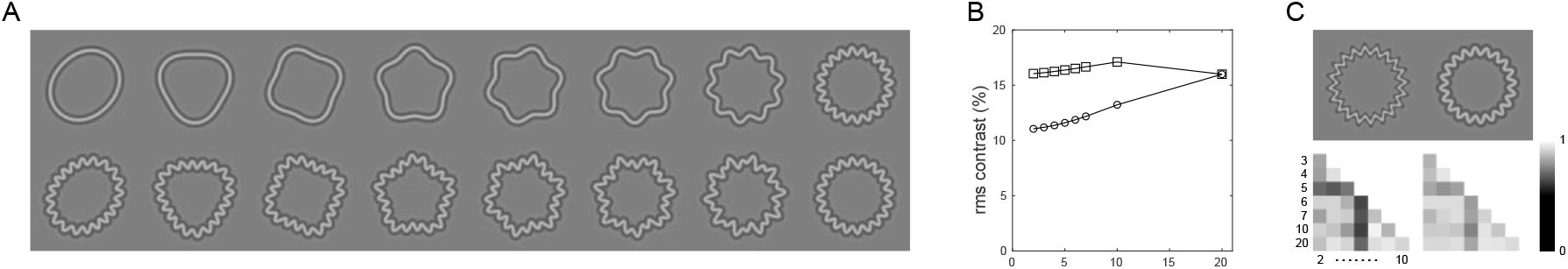
The radial frequency (RF) stimuli and their contrast and spatial frequency properties. A: RF stimuli presented to participants (top row) and their RF-ripple counterparts (bottom row). Exemplars are shown for a single orientation only. During the acquisition of BOLD responses, however, six orientations of each of the stimuli shown were presented. B: The root-mean-square (rms) contrast of the stimuli computed as the mean over the annulus that capture all contrast variations of the six orientations presented. Open circles and squares are for the RF and RF-ripple stimuli. C: The top two panels show different rendering approaches; the luminance profile is computed on the basis of the radial distance from the contour (left) or the perpendicular distance from the contour (right). The rendering techniques result in different spatial frequency variations between our stimuli as shown in the correlation between the two-dimensional amplitude spectra of stimuli at different radial frequencies (lower two panels). Moreover, radial rendering produces lower and more varied correlation between spatial frequency content of RF stimuli (bottom left) than the rendering derived from the perpendicular distance that was used in the present study (bottom right).

We also took a precaution to largely equate our stimuli for spatial frequency content. Spatial frequency content can change with amplitude and radial frequency when the contrast profile of the stimulus is computed from the radial distance from the RF contour alone (Salmela et al., 2016). We used the distance perpendicular from the RF contour as the input to the fourth derivative function that defined the contrast profile of our stimuli. Our approach has the effect of all but removing and variation in spatial frequency content (Figure 2C). The alternative rendering of the profile uses only the radial distance from the contour to compute the fourth derivative, which is entirely appropriate at threshold, but does introduce greater variability in spatial frequency content for suprathreshold stimuli that we use (Figure 2C).

The approaches we took to control for contrast and spatial frequency content of stimuli allow us to investigate responses that are related to low radial frequencies that are known to be processed globally and differentiate them from responses that are largely driven by variations in contrast energy and spatial frequency (Salmela et al., 2016). It should also be noted that orientation content of the stimuli scales with radial frequency of the RF stimuli in much the same way as contrast energy but will again be largely equated in the RF-ripple stimuli, so we will refer to contrast energy and orientation content together when discussing the results.

#### Experimental design

Each radial frequency tuning scan started with a 10s fixation period, followed by two cycles of stimulus blocks. Each block lasted 6s, during which each RF pattern was repeated six times (0.8s presentation and 0.2s interstimulus interval; phase of each pattern was randomly selected between the range of 0:60:300 degrees). The order in which the stimuli were presented followed the design used by Harvey et al., (2013): a ramp-up sequence, with RF2-7 and RF10, followed by 24s of RF20 (‘baseline’), and a ramp-down sequence (RF10, RF7-2) followed by another 24s of RF20. Each scan terminated with 20s of fixation to capture the full hemodynamic response for the final stimulus. The same procedure was used for both standard RF stimuli and RF-ripple stimuli. Participants performed the same oddball (contrast reversal) task as described for the stimulus localiser, to ensure attention was maintained.

#### Modelling

The data were first pre-processed using FEAT: the first 5 volumes were removed to allow magnetisation to reach a steady state, followed by high-pass filtering (cutoff 100s), slice timing correction and motion correction (MCFLIRT). No spatial smoothing was applied to the data, and all runs were coregistered to each participant’s high-resolution structural space. We then extracted and concatenated the timeseries of all voxels from the restricted ROIs. Additionally, 20,000 randomly selected white matter voxels timeseries were extracted to create a null distribution for modelling.

To model the data, we applied a gaussian function to our experimental design, parameterised by a *mean* preferred RF plus a *tuning width* (SD). This was convolved with a standard double gamma hemodynamic response function (hrf) to estimate the BOLD response we would expect based on the respective tuning parameters. Note that we did also test logarithmic gaussian models, but no improvements were found.

This gaussian predictor was concatenated for all runs, we also included separate run-wise predictors for the temporal derivative (to allow slight temporal deviations across runs), a constant, a predictor for oddball events (again convolved with default hrf) and motion confound covariates. All run-wise predictors were set to zero outside their respective runs, and all predictors were high-pass filtered to match the data. After fitting these predictors to our data, we took the sum of squared residuals as our estimate of model accuracy.

To perform the fitting, for each voxel we first tested approximately 3,500 initial models, with means ranging −2.5:20 (increments of 0.25) and tuning widths (SDs) ranging 0.5:20 (0.5 increments). The best fit for each voxel was then further refined using non-linear least squares optimisation (Matlab’s *lsqnonlin*). The fitting limits were set to −5 ≤ Mean ≤ 30 and 0 < SD ≤ 30; constraining the mean preferred RF to be reasonably close to our stimulus range to enforce plausible fits.

The white matter voxel fits were used to generate a null distribution of fit accuracy and from this we calculated (two-tailed) significance for our fitted ROI voxels. We only kept ROI voxels that remained significant after correction for multiple comparisons (accounting for all included ROI voxels using Benjamin & Hochberg FDR correction). We also excluded voxels whose fitted parameters were within 0.1 units of our fitting limits (i.e. Mean ≤ −4.9 or ≥ 29.9; SD ≤ 0.1 or ≥ 29.9), as we may not have found the best possible fit for such voxels and even if we had, the interpretability of fits so close to the limits would be questionable.

To assess the distribution of the fits across participants, we nonparametrically estimated the probability density functions (PDFs) of each ROI using kernel density estimation (*ksdensity* in Matlab; normal kernel, default bandwidth, 200×200 equally spaced points encompassing fitting limits). We then graphically inspected the resulting distributions for each ROI via density plots. We pre-empt those results here to allow a fuller description of the methods we used to assess them quantitatively (below): Two clusters emerged (see Figures 2 and 3) tuned to low and high radial frequencies.

**Figure 3.**
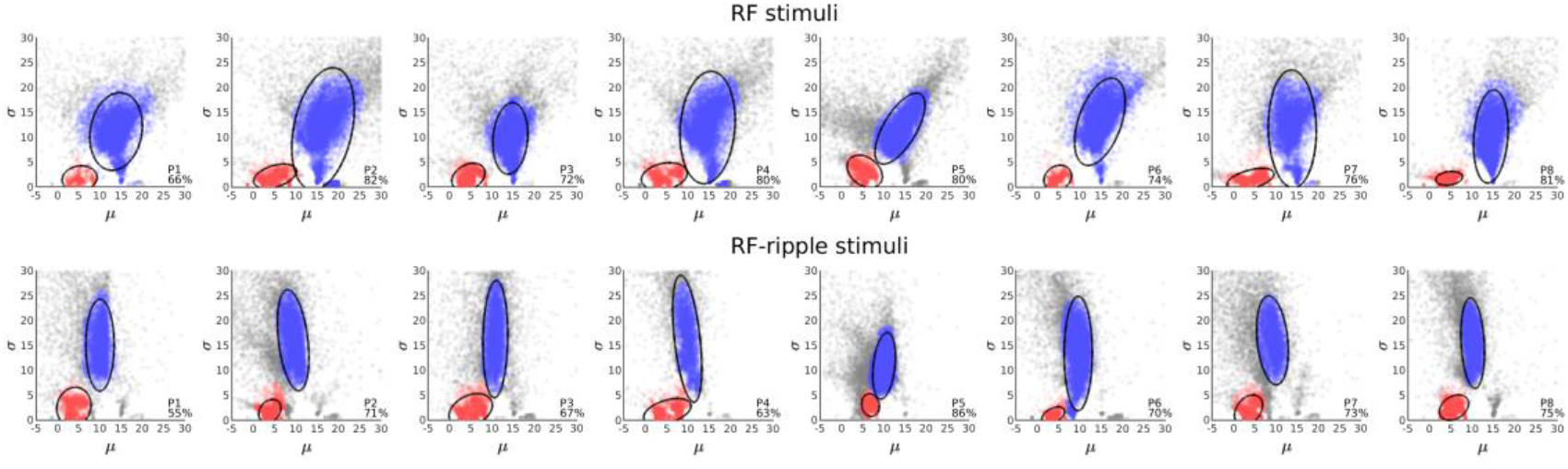
Model fits for each individual. Each panel plots the radial frequency tuning center, μ, and bandwidth, σ, for every model that exceeding statistical thresholding. The top and bottom rows shows data for the RF and RF-ripple stimuli, respectively. The model parameters fall into two clusters shown in blue and red. Tuning to low radial frequency, with small bandwidths (red) was evident in each participant and was largely unchanged for the stimulus set. Tuning to higher radial frequency was also common to all participants (blue) but differed for the different stimulus sets. Inset in each panel is the percentage of the voxels identified in localizer scans that survived thresholding.

To further quantify and characterise the properties of the fits, we performed a whole-brain (whole FOV) clustering analysis using density-based spatial clustering (*dbscan* in Matlab). Both preferred mean (RF tuning) and SD (tuning width) were first standardised (z-scores) to ensure neither dominated the clustering, and we specified a minimum of 500 voxels to form a cluster. The search radius (epsilon) was first estimated using a k-distance graph, then manually adjusted to best segment the observed clusters, while including as many voxels as possible. The preferred RF and corresponding tuning widths were then extracted for each resulting cluster. To inform upon the distribution of radial frequency tuning within each ROI, we calculated the proportion of each ROI’s voxels that fell into a given cluster obtained from the whole-brain analysis approach.

#### Representational Similarity Analysis

Similarity of neural representations between radial frequency patterns in each ROI and across stimulus sets was tested using Multivoxel Pattern Activity analysis. Specifically, voxel-wise patterns of activation for each session, ROI, and participant were extracted by first concatenating all scans within one session, and then building a corresponding GLM model. Here, each RF pattern, apart from RF20, which was treated as ‘baseline’ common to both sessions, was entered as a regressor of interest. The relative temporal derivatives, together with confound regressors modelling the oddball events in each scan and the six FSL-generated motion regressors were added to the model. After extracting activity patterns, we ran Representational Similarity Analysis (RSA) by correlating the voxel-wise activity patterns for all pairwise combinations of RF patterns for each ROI and stimulus set. We then established predicted similarity matrices on the basis of radial frequency differences between items in each stimulus set. Two predictors were used; one based on the log ratio of stimulus item pairs and the other on the difference in frequency of the pairs. The former prediction is a better approximation to how radial frequency is perceived - unit differences between low radial frequencies such as 2 and 3 are perceived as far greater than the same differences between high radial frequencies such as 19 and 20.

### Statistical Analysis

The design of the study is repeated measures with a relatively small number of participants examined in detail. We implement inferential statistical tests – most frequently ANOVAs - to examine effects of stimulus set and region of interest on outcome measures. In the cases where interactions were observed we followed up with appropriate post hoc F and T tests to examine specific effects. Data met assumptions for the tests, although in sometimes only after appropriate transformation of the data to ensure equal variance assumptions were met. The raw and transformed data are presented in the manuscript. We also used correlation as a measure of relationships between variables and subjected them to inferential tests after appropriate Fisher transformation of the data to ensure normality. With a relatively small number of participants the statistical tests used reached significance only when the vast majority, if not all, participants exhibited an effect, consistent with type one errors being appropriately minimised.

## Results

To evaluate the ventral pathway’s responses to and representation of radial frequency patterns we conducted the following analyses and report on them under different heading within the Results section. First, we examined the *distribution of radial frequency tuning and bandwidths* across voxels that fell within all the ROIs we identified, and then separately for each ROI. Second, we performed a *quantitative analysis of low and high radial frequency tuning clusters* that were evident in the distributions. This allowed us to examine the effects of two stimulus sets - RF and RF-ripple - and ROI on the tuning to radial frequency. Third, we examined the *representation of radial frequency in the ventral pathway* using representational similarity analysis.

### The distribution of radial frequency tuning and bandwidths

Our first assessment of the radial frequency tuning properties was to scatter plot (Figure 3) model parameters – the mean, μ, and standard deviation, σ, - for each participant collapsed over all regions of interest. This serves two purposes; first, it captures what the model characteristics are and whether they cluster and second whether they vary across participants or between the stimulus sets (RF-stimuli and RF-ripple stimuli). The scatter plots show that models that survive statistical thresholding fall into two clusters in each participant, one in the lower left of each plot, (in red) corresponding to narrow tuning to low radial frequencies, which are globally processed, and the other located more to the right in the plot (in blue) reflecting broad tuning to higher radial frequencies. The location of the clusters in the scatter plots vary little across participants. Furthermore, the addition of the ripple to the stimuli shifts the cluster centred on high radial frequencies to a lower radial frequency, but the low radial frequency tuned cluster appears largely unchanged. Because we have assessed responses from a relatively small number of participants in detail, it is important to demonstrate that the general characteristics of the signals we record are shared across all participants (as shown in Figure 3). This feature has a strong bearing on inferential statistics that we present later, which are only likely to show significant effects if all participants of a small group share very similar response characteristics.

The ellipses shown in the scatter plots capture 95% of the models in each cluster and we use the co-ordinates of centre of the ellipses as a measure of the cluster’s central tendency. We also captured the proportion of voxels in the ROIs that survived thresholding (inset in each panel of Figure 3). The values are relatively high (55-86%) given that the ROIs were defined by high powered block design experiments that captured responses to the annular region (compared to uniform gray) of the visual field where stimuli were presented in contrast to the stimulus regime used to capture radial frequency tuning in which the target stimuli were presented for the vast majority of the time.

The data shown in Figure 3 are aggregated in a two-dimensional histogram of the radial frequency tuning and bandwidths (Fig 4 A and C). As expected from the individual data, two clusters are evident, one in the lower left of the histograms which corresponds to a narrowband tuning at low radial frequencies, and the other more centrally located in the histogram and corresponding to broadband tuning to higher radial frequencies. Superimposed on the histogram are the cluster centres for each participant, which demonstrate narrow dispersion of this measure between participants.

**Figure 4.**
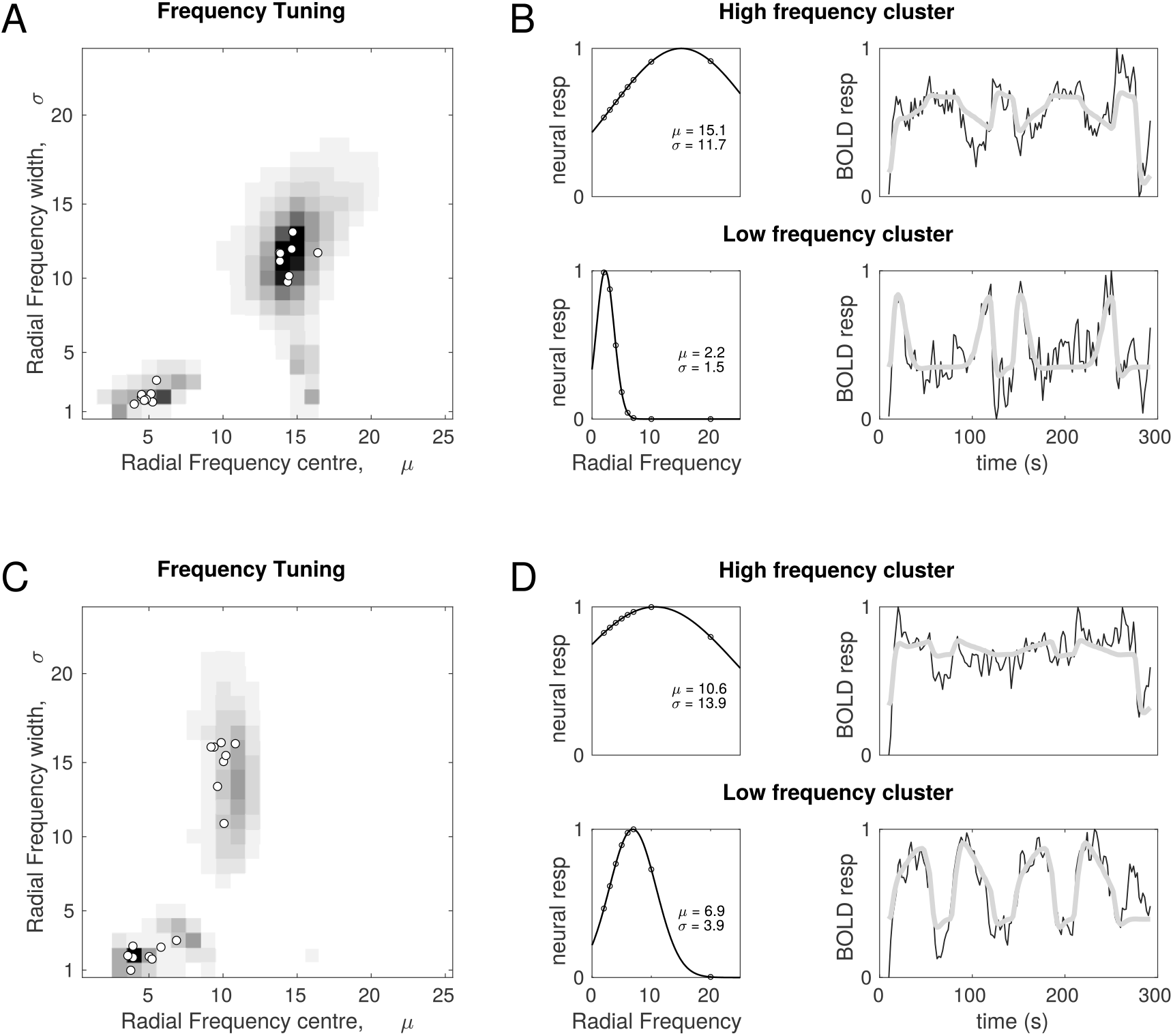
Radial frequency tuning properties of visual cortex. Panels A and C are grayscale renderings of a two-dimensional histogram of the modelled radial frequency tuning of voxels in all regions of interest (V1-V4, LO-1, LO-2 and LO) for the RF and RF-ripple stimuli, respectively. Modelled tunings were retained only if significant, which was the case for 70% (RF) and 68% (RF-ripple) of voxels in the stimulus representation. Tunings that are representative of the clusters found at low and high radial frequency are shown in panels B and D for the RF and RF-ripple stimuli, respectively. The population neural response as a function of radial frequency are shown to the left with the upper and lower graphs depicting high and low frequency cluster tunings. On the right examples of time series data (black) and the best fitting model (thick grey) derived from the population neural responses shown in the adjacent graphs are shown. The top rows of B and D show responses that capture contrast and orientation content of the stimuli as shown in Figure 2, with a largely monotonic increase in response as a function of radial frequency for the RF stimuli, but not for the RF-ripple stimuli. The bottom rows of B and D show tunings to different radial frequencies, 2.2 and 6.9, to illustrate the range of tunings observed in the low frequency cluster common to both the RF and RF-ripple stimuli.

Examples of the radial frequency tuning functions and the way that they model the BOLD time series are shown in Figure 4B and D. For the RF stimuli the representative tuning to the high radial frequency (μ = 15.1), which is also relatively broadband (σ = 11.7), exhibits a broadly monotonic increase in response with increasing radial frequency (left upper graph Figure 3B) similar to the contrast energy of the stimuli that are shown in Figure 2B. For the RF-ripple stimuli (left upper graph Figure 3D), the representative tuning to high radial frequency is centred on a lower radial frequency (μ= 10.6) and has a higher bandwidth (σ = 13.9) than for the RF stimuli. The tuning function (left upper graph Figure 3D) no longer has the monotonic relationship between radial frequency and there are smaller variations in response as a function of radial frequency overall, again capturing quite well the relationship between contrast energy and radial frequency of the RF-ripple stimuli shown in Figure 2B. The tunings to high radial frequency appear therefore to be common and are consistent with responses driven by contrast and likely orientation rather than global shape properties.

The tunings to low radial frequencies (lower left graphs of Figures 4B and D) do not exhibit response profiles that fit variations in contrast energy or orientation content of the stimuli for either the standard RF of RF-ripple stimuli. Moreover, the highest contrast stimuli are non-preferred stimuli for these tuning functions. As a result, the best fitting models derived from the tunings to low radial frequency differ markedly from those derived from the high radial frequency tunings (graphs to the right in the upper and lower rows of Fig 4 B and D). The low frequency tuning clusters for both the RF and RF-ripple stimuli cover the range of radial frequencies that are processed globally; circa 2-7. We show selected time series and model fits for radial frequencies near the limits of this range, namely 2.2 in Figure 3B and 6.9 in Figure 3D to capture how the time series reflect different (2.2 vs 6.9) low frequency tunings.

The next step was to assess the tuning preferences of each ROI (averaging across participants). We did this by computing probability density functions (PDFs) using kernel density estimation. This approach is broadly analogous to a smoothed 2D histogram and makes no assumptions about the underlying distribution of the data, allowing for further examination of the radial frequency tuning distributions for different regions of interest and for the RF and RF-ripple stimuli (Figure 5). Predictably this approach revealed the same two general tuning clusters that were detected when data were collapsed across regions of interest (Figures 3&4). However, the high and low frequency tuning clusters occurred to different extents in different regions of interest. The high frequency tuning dominated in early visual cortex (V1-3), while the low frequency tuning became increasingly prominent up the visual hierarchy and dominated in LO. There were also understandable differences between the stimulus conditions. For the RF-ripple stimuli the tuning to high frequencies was less common overall and largely absent in LO-2 and LO. In all regions exhibiting the high frequency tuning it was also centred on lower frequency for the RF-ripple than the RF stimuli as previously noted. The tuning to low radial frequencies appears less affected by the change from RF to RF-ripple stimuli and is therefore more consistent with mechanisms processing global shape properties.

**Figure 5.**
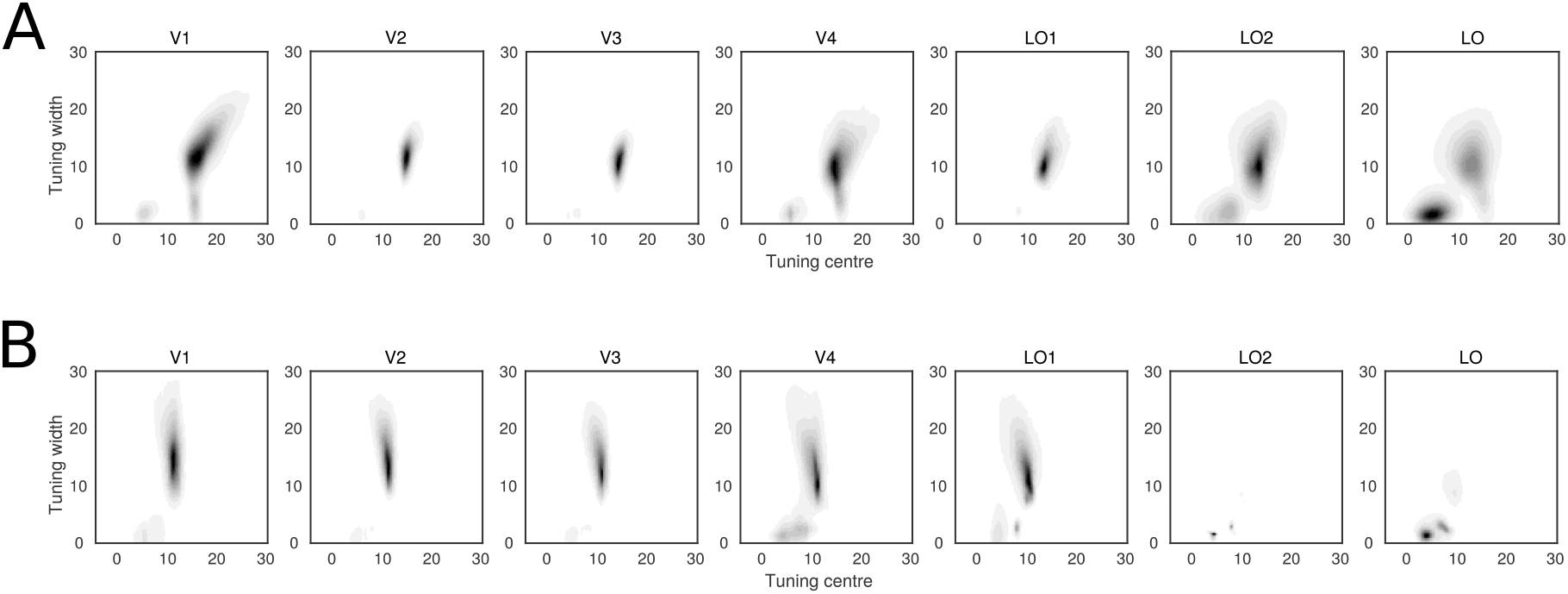
Radial Frequency tuning for regions of interest. The model fitted parameters, preferred radial frequency tuning and tuning width, mapped onto two dimensions. The grayscale maps the relative number of model fits - darker colours represent more voxels. Two clusters of model fits emerged, one with broad tuning to high radial frequency that dominated model fits in early visual areas and another narrowly tuned to low radial frequency that emerge in V4 and are prominent in lateral occipital areas.

### Quantitative analysis of low and high radial frequency tuning clusters

The visualisation of the data above was for the mean data across all participants. To isolate the two clusters for each individual, we took all fitted voxels (initially agnostic of ROI) per stimulus set and participant and clustered them. This was straightforward for RF stimuli as the two clusters were well separated, while clusters were separated less for the RF-ripple stimuli, nonetheless, the clustering results showed notable consistency across participants (as demonstrated by 95% CIs below).

We found the ‘high-RF’ cluster shifted and broadened across stimulus conditions as could be seen in Figures 3&4. Specifically, the mean preferred radial frequencies in the RF and RF-ripple experiments were 14.50 and 9.90 (95% CIs: 13.92-15.09; 9.95-10.26 respectively), and tuning widths were 11.41 and 14.94 (95% CIs: 10.68-12.15; 13.62-16.26 respectively). This cluster appears to encompass voxels that have no specific preference to radial frequency, with a bias towards higher frequency stimuli, likely due to their greater contrast energy and orientation content. As such, the leftward shift and broadened tunings for RF-ripple versus standard RF stimuli make sense; the RF-ripple stimuli had higher contrast energy and orientation content (introduced by the RF 20 ripple) that varied less across the stimulus set, and so voxels that previously preferred e.g. RF20 would now be expected to respond more equally across the stimulus range.

The second ‘low-RF’ cluster was almost identical across stimulus sets; the mean preferred radial frequency for the RF and RF-ripple experiments were 4.80 and 4.76 (95% CIs: 4.46-5.13; 3.95-5.57 respectively), and tuning widths were 2.03 and 2.08 (95% CIs: 1.68-2.37; 1.65-2.51 respectively). The comparable tunings across both sessions suggests that there is a subset of voxels that are sensitive to shape, defined by radial frequency changes, in a more global sense, rather than sensitivity to local contour information alone. Given that this cluster analysis was performed over all ROIs, we further mapped the clusters to the ROIs to determine where the voxels that had a preference for the low and high RFs could be found.

To map the two clusters (low-RF, high-RF) from each stimulus set to individual ROIs, we calculated the proportion of each ROI’s voxels that fell into a given cluster. We assessed proportions for both clusters as voxels could be outside the two clusters (i.e. classed as ‘noise’ – see gray data points in the scatter plots shown in Figure 3), so if an ROI had more voxels in one cluster it did not necessarily imply a proportional decrease in the other. The proportions of voxels falling into the low and high clusters are bar charted in Figure 6A and B for the RF and RF-ripple stimuli, respectively. In both charts the proportion of voxels showing tuning to low frequencies increases up the visual hierarchy and in LO dominates for the RF-ripple stimuli.

**Figure 6.**
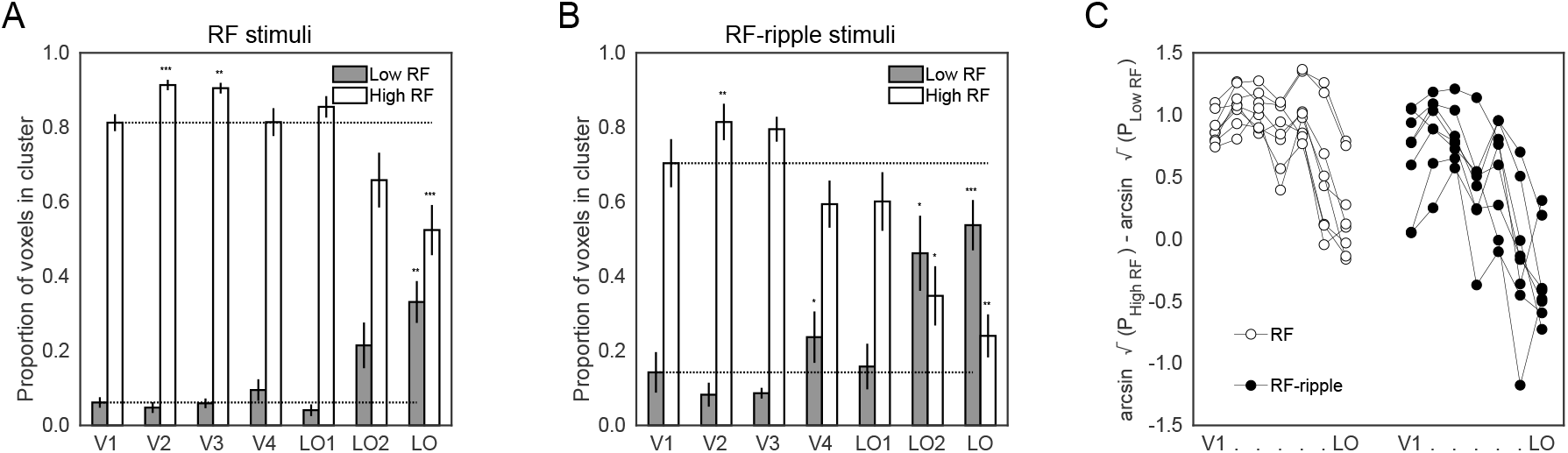
The proportion of model fits that fell into high (light bars) and low (dark bars) radial frequency clusters. A: Data are shown for the RF stimuli. B: Equivalent data for the RF-ripple stimuli. Significant differences from V1 proportions are denoted by asterisks as follows; *<0.05, **<0.01, ***<0.001 as computed from planned contrasts (see text for details). C: The data in A and B were submitted to a three-way ANOVA after transforming the data to ensure they met the assumptions of equal variance. Transformed data are plotted to capture the difference between voxels tuned to high and low radial frequencies for each participant across all ROIs and the two stimulus sets, which underpins the three-way interaction reported in the text.

The proportions were arcsine transformed (sin^−1^√p) to meet normality requirement for a 2×2×7 (Stimulus x Cluster x ROI) repeated measures ANOVA, which was applied to the data. This resulted in a significant three-way interaction (F(6,42) = 5.51, *p* = 2.8×10^−4^), which we help illustrate in Figure 6C, where the difference between the transformed proportion of voxels tuned to high and low frequencies is given for each individual for each ROI. Evident in each participant’s data in the plot is the reduced difference between high and low frequency tuned voxels up the visual hierarchy and even greater reduction in this difference for the RF-ripple stimuli. Subsequent ‘Cluster x ROI’ ANOVAs also yielded significant interactions between the two factors (for both stimulus sets *p* < .001). To follow up on the interactions we ran 4 one-way repeated measures ANOVAs exploring the main effects of ROI separately per stimulus condition and cluster.

ROI had a significant effect on the proportion of voxels mapped onto each cluster tuned to high or low radial frequency patterns for both RF and RF-ripple stimuli (RF high/low: F(2.20,15.37) = 24.12, *p* = 1.3×10^−5^; F(2.25,15.77) = 12.90, *p* = 3.5×10^−4^, respectively; RF- ripple high/low: F(2.44,17.11) = 23.98, *p* = 5.1×10^−6^; F(1.80,12.59) = 13.54, *p* = .001, respectively). While this was informative, suggesting that it is possible to differentiate a preference in processing local (high radial frequency) versus global (low radial frequency) shape information, it did not allow us to identify where radial frequency tuning preferences began to diverge from low-level representations. To this end we ran planned comparisons, with V1 as ‘baseline’ where, on average 64.63% of voxels were in the high radial frequency cluster, compared to 13.67% in the low radial frequency cluster when the RF stimuli set was used. Similarly, in the RF-ripple session 57.83% and 19.85% of voxels were mapped to the high and low radial frequency clusters, respectively.

First for the high radial cluster, in RF experiment we found that V2 and V3 had significantly more high radial frequency tuned voxels compared to V1 (V2: 73.35%, p = 3.8×10^−4^; V3: 72.47%, *p* = .005) whereas only LO had significantly fewer high radial frequency tuned voxels (46.61%, *p* = 2.9×10^−4^). In the RF-ripple experiment, both LO2 and LO had significantly fewer high radial frequency tuned voxels (LO2: 42.57%, *p* = .012; LO: 47.11%, *p* = .001), only V2 had significantly more (65.58%, *p* = .007). No other results were significant (all *p* > .090).

For the low radial frequency cluster, LO had significantly more low radial frequency tuned voxels compared to V1 across both experiments (RF experiment: 34.36%, *p* = .001; RF-ripple experiment: 47.11%, *p* = 1.4×10^−4^). In addition, in the RF-ripple experiment we also found greater proportions of low radial frequency tuned voxels in V4 and LO2 (V4: 27.57%, *p* = .019; LO2: 42.57%, *p* = .024). No other results were significant (all *p* > .062).

In summary, as highlighted earlier, tunings for high radial frequencies dominate in early visual cortex (V1-V3). Conversely, LO consistently diverged from V1, by showing the greatest preference for low radial frequency patterns. However, V4 and LO2 did show differences from V1 for the RF-ripple experiment when variations in low-level contrast and orientation content in the stimuli were much reduced.

### Representation of radial frequency in the ventral pathway

One consequence of LO’s low radial frequency tuning preference is that we would expect its pattern of activity to show consistency across the RF and RF-ripple stimuli, as both sets of stimuli share comparable global profiles. Conversely, for an ROI such as V1, which is likely responding to the contrast energy and orientation of our stimuli we would expect different patterns of activation for the two sets of stimuli.

To test this, we used MVPA to extract the voxel-wise patterns of activation related to each stimulus pattern for each stimulus set, ROI and participant. Representational Similarity Analysis (RSA) was then computed by correlating the voxel-wise activity patterns for all pairwise combinations of radial frequency patterns (for each ROI per stimulus set). This generated similarity matrices, capturing neural similarity across radial frequency patterns, which are shown in Figure 6A&B for V1 and lateral occipital ROIs. For the RF stimulus set the similarity in patterns of activation are relatively high across all regions. For the RF-ripple stimulus set large changes in similarity are observed for V1 (and other ROIs - V1-V4). In lateral occipital ROIs similarity is better preserved from LO-1 through to LO. In LO, for example, the similarity matrix for RF and RF-ripple stimuli is largely unchanged (Figure 7A and B).

**Figure 7.**
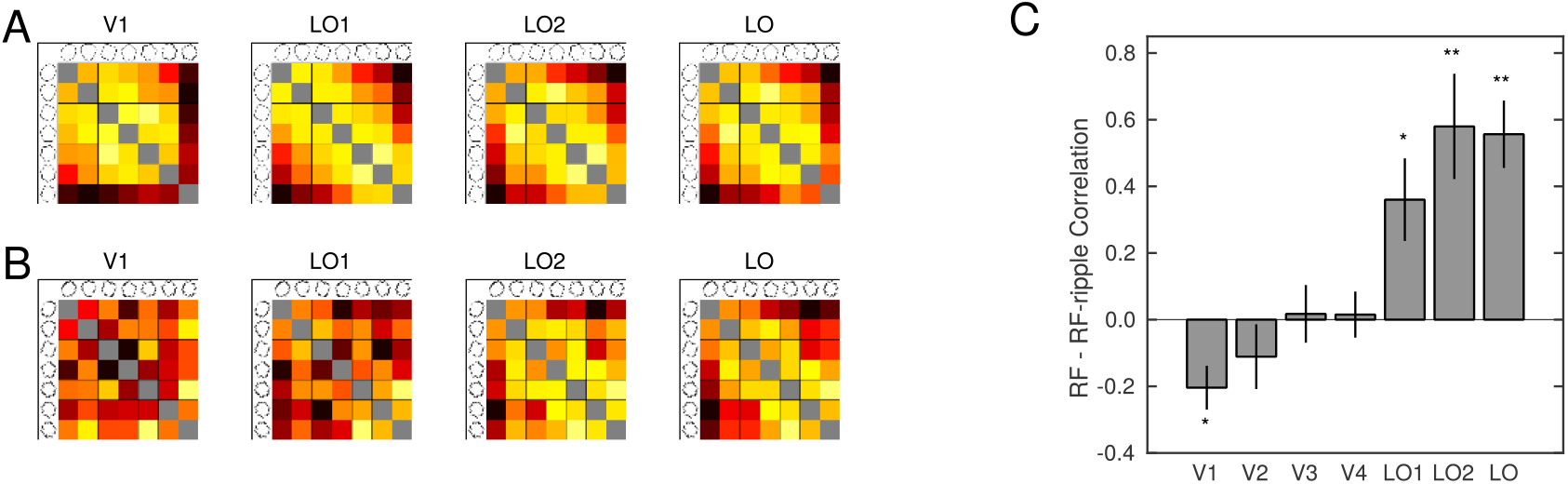
Similarity of responses to RF patterns across visual cortex. **A**: Matrices of correlation between responses to RF patterns displayed as heat maps. Data are shown for ROIs V1, LO-1, LO-2 and LO with the top and bottom rows showing correlations for the RF and RF-ripple stimuli, respectively. **B**: The correlation between the similarity matrices obtained from responses to RF (top row matrices) and RF-ripple (bottom row matrices) stimuli for each ROI. Asterisks denote the following; * p < 0.05, ** p < 0.01.

To test statistically how well the similarity was preserved across stimulus conditions, for each participant’s ROI we correlated that ROI’s similarity matrix for the RF stimuli (Fig 7A) with its corresponding RF-ripple similarity matrix (Fig 7B) and ran one-sample t-tests on the (Fisher-Z transformed) results (Fig 7C). We found V1 had a significant negative correlation (t(7) = −3.55, *p* = .009), but only LO1, LO2 and LO showed significant positive correlations (t(7) = 2.61, 4.27, 5.98, *p* = .035, .004, .001 respectively). No other results were significant (all *p* > .305).

To test whether the relationships between patterns of response to radial frequency stimuli were indeed related to differences in the radial frequency of the stimuli we presented, we created two predictor similarity matrices. The first used the log ratio of the radial frequency of pairs of RF patterns (e.g. for RF2 and RF3; -abs(ln(2/3)) = −0.41). We used log ratios as we reasoned that the jump from e.g. RF2 to RF3 perceptibly much greater than the jump from e.g. RF20 to RF21, and so the similarity matrix should capture this. The second predictor matrix simply computed the difference in radial frequency of the stimuli. This predictor might better capture variations in contrast and orientation, which were relatively large for the RF stimulus set and scaled with radial frequency. The predictor similarity matrices were then correlated with all neural similarity matrices across ROI and stimuli (Figure 8).

**Figure 8.**
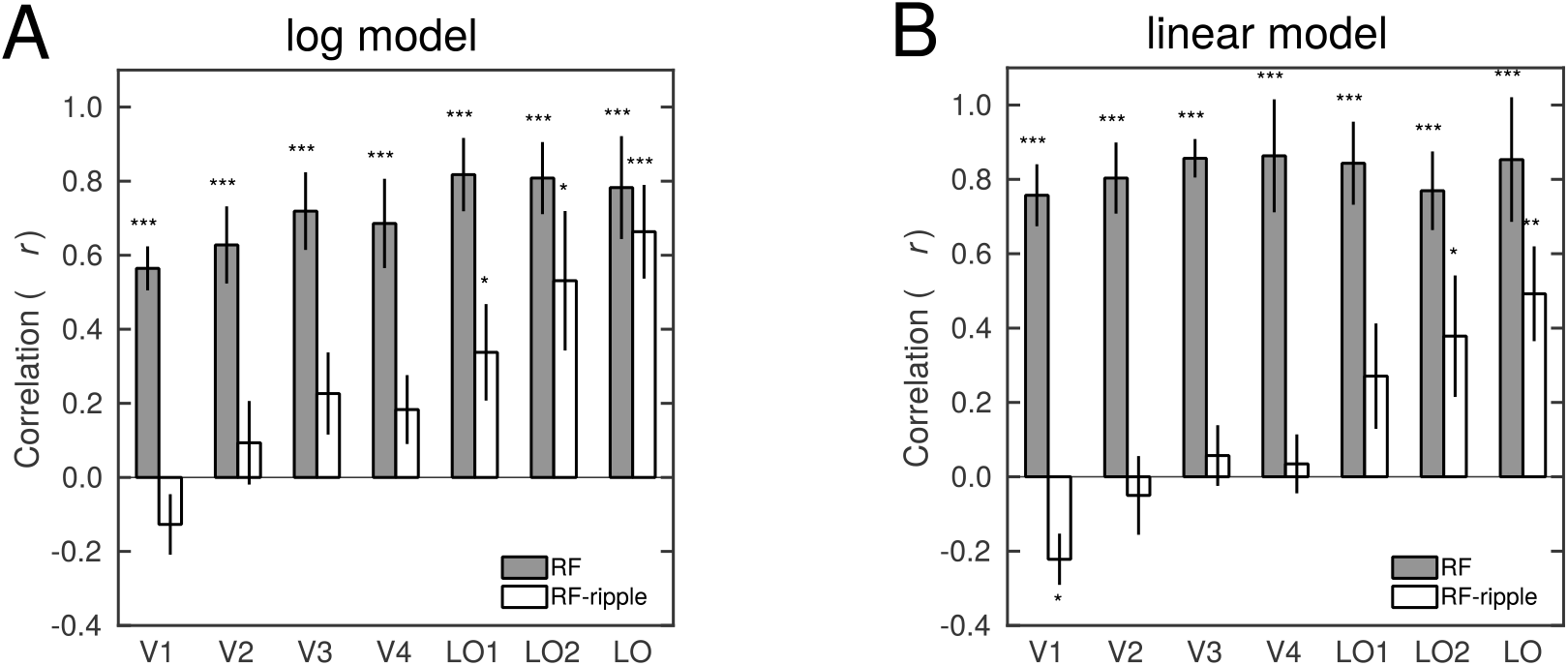
Stimulus similarity predicts response similarity. Correlations are given between the response similarity matrices (as shown in Figure 7) and a similarity matrix constructed from the log difference between the radial frequency of stimuli. Data are given for the two stimulus sets, RF and RF-ripple, shown by grey and white bars, respectively. *, ** and *** denote that the distributions of the correlations are significantly different from zero at p < 0.05, 0.01 and 0.001, respectively.

Data for the log model predictions are given in Figure 8A and the extent to which the model predicts the similarity of neural responses was assessed by computing single sample t-tests. The similarity of neural responses is predicted very well by the log model for the RF stimulus set (p < .001 for all ROIs), but for RF-ripple stimuli the log model captures similarity of the neural responses only in LO1, LO2 and LO (t(7) = 2.685, 3.10, 6.29, *p* = .031, .017, <.001, respectively). No other results were significant (all *p* > .077). Data for the linear model are presented in Figure 8B and were assessed in the same way. Again, the similarity of neural responses to RF stimuli were predicted well in all ROIs (p < .001), while for the RF-ripple stimuli the linear model significantly captured the similarity of neural responses only in LO2 and LO (t(7) = 2.42, 4.21, p = .046, .004, respectively). There was also a significant negative correlation between the linear model’s predictions and the similarity of neural responses in V1. No other results were significant (all *p* > .092). While the alternative models appear to have similar predictive power, it is noteworthy that the linear model has higher correlation than the log model with similarity of neural response to the RF stimuli. In contrast, for the RF-ripple stimuli it is the log model that performs better, at least for some regions of interest.

We assessed the two models’ predictive power as a function of stimulus set and region of interest by performing a three-way ANOVA of the Fisher-transformed correlation data. We acknowledge that caution must be taken in interpreting such an analysis particularly for regions of interest in which the models have low correlations with the similarity of neural responses. The three-way ANOVA showed a three-way interaction (*F* = 4.194, *p* = .002, *η^2^* = .375) meaning that the extent to which the models fitted the data varied with stimulus type in a different way across regions of interest. Also evident was the significant two-way interaction between the model and the stimulus type (*F* = 39.583, *p* < .001, *η^2^* = .850) likely driven by the observation that the linear model worked best for the RF stimuli while the log model accounted better for pattern of responses to the RF-ripple stimuli.

In Table 1 we show the results of the follow-up analysis in which we ran two-way ANOVAs in each ROI and paired t-tests. Two-way interactions were evident in V1 to V4 and LO. In V1 and V2 the interaction was driven by a better fit of the linear than the log model for the RF stimuli and an absence of differences in model fit for the RF-ripple stimuli, where both models performed very poorly. In V3 and V4 a similar advantage of the linear model was evident for the RF stimuli, but now the log model worked better than the linear model for the RF-ripple stimuli. While it is tempting to take the latter at face value, the correlations are low and straddle zero across participants, so even though statistically significant this increase in model fit may not be meaningful. In LO the interaction was driven by significantly better performance of the log model over the linear model for RF-ripple stimuli and largely similar performance of the models for RF stimuli. This pattern of model fits is the opposite of that observed in V1. It is also important that in LO all model fit correlations were significantly different from zero allowing the improved fit of one model over the other to be taken as meaningful.

**Table 1:** Results of two-way ANOVA applied to each region of interest. Region of interests are given in each row and columns show data for the interaction (model and stimulus), main effects of model (linear vs log) and stimulus (RF vs RF-ripple), and post-hoc T-tests for differences in the models for each stimulus type.

| **ROI** | **Model x Stim**<br>**F, p, $\eta^2$** | **Model**<br>**F, p, $\eta^2$** | **Stim**<br>**F, p, $\eta^2$** | **RF**<br>**lin - log**<br>**T, p** | **RF-ripple**<br>**lin - log**<br>**T, p** |
| --- | --- | --- | --- | --- | --- |
| **V1** | **13.906, .007, .665** | **56.539, <.001,**<br>**.890** | **191.976, <.001,**<br>**.965** | **4.932, .002** | -1.870, .104 |
| **V2** | **13.877, .007, .665** | **10.338, .015, .596** | **100.267, <.001,**<br>**.935** | **4.455, .003** | -2.103, .084 |
| **V3** | **38.626, <.001,**<br>**.847** | 3.098, .122, .307 | **88.049, <.001,**<br>**.926** | **4.237, .004** | **-3.377,**<br>**.012** |
| **V4** | **44.253, <.001,**<br>**.863** | 10.414, .015, .598 | **31.541, .001, .818** | **5.838, .001** | **-2.894,**<br>**.023** |
| **LO1** | 3.389, .108, .326 | .002, .965, .000 | **50.855, <.001,**<br>**.879** | .602, .566 | -.901, .398 |
| **LO2** | 2.006, .200, .223 | 1.970, .203, .220 | **16.915, .005, .707** | -.828, .435 | -2.062, .078 |
| **LO** | **8.540, .022, .550** | .066, .804, .009 | **8.552, .022, .550** | 1.459, .188 | **-3.445,**<br>**.011** |

Overall therefore, the MVPA results largely mirror the univariate findings; highlighting a move away from local towards more global or holistic processing that becomes clear and likely dominant in the Lateral Occipital cortex. Global processing of shape also appears in retinotopically defined LO1 and LO2. Together the results reflect a processing pathway of shapes that moves from a low-level retinotopic representation towards a more abstract representation in anterior Lateral Occipital cortex.

## Discussion

We found two characteristic tuning profiles to radial frequency stimuli. The tuning most commonly detected in the visual cortex was broad band and centred at relatively high radial frequencies for which there is no behavioural evidence of global processing. Our stimulus manipulation that added a high radial frequency component to the radial frequency patterns shifted the tuning’s centre indicating that low level stimulus features that include contrast and orientation content were driving this aspect of the cortical response. The second tuning we detected was centred on the low radial frequencies, for which there is strong behavioural evidence of global processing (Hess et al., 1999; Jeffrey et al., 2002; Loffler et al., 2003), and had a narrow bandwidth. This tuning remained, when the high radial frequency ripple was added to our stimuli, which we, in common with others (Bell et al., 2007a), take as an indicator that the global properties of the shape were driving this aspect of the cortical response. We found that the low radial frequency tuned responses were most evident in the extrastriate cortex, particularly in LO, but the high frequency ripple added to stimuli - to render them less varied in orientation and contrast content - highlighted that this tuning to global shape could also be revealed in retinotopically defined areas LO1, LO2 and V4. Multivariate approaches showed that all the subdivisions of visual cortex we examined exhibited a relatively high level of similarity in the patterns of response across radial frequencies, but that this pattern was disrupted very much in primary visual cortex and other early retinotopic areas, when the high radial frequency ripple was added to stimuli. Indeed, the only areas that maintained the similarity across radial frequency representations once the ripple was added were LO1, LO2 and LO. Finally, the similarity of the stimuli as quantified by the difference of the log radial frequency provided a good prediction of how the pattern of responses were similar, consistent with human sensitivity to the global shape of radial frequency stimuli.

### Stimulus properties that drive responses in early visual cortex

The responses of early visual cortex to RF stimuli revealed a strong bias to stimuli that exhibited the greatest contrast and orientation content. Given that contrast and orientation increased systematically with increasing radial frequency of the RF stimulus set, the bias to contrast and orientation was expressed as a tuning of responses to high radial frequency. This tuning was prevalent and dominated in early visual cortex. Our findings are consistent with early visual cortex having the most expansive contrast response function (Gouws et al., 2014) and well-known orientation selective neurons (Hubel & Wiesel, 1968). Adding a high radial frequency component that defined the RF-ripple stimuli had largely predictable effects on the tuning we measured from early visual areas. Because the addition of the ripple reduced the contrast and orientation variations across the stimuli and also changed the relationship of those variations with radial frequency, the high radial frequency tunings in early visual cortex shifted to a lower, although still remaining high, radial frequency. The results fit well with the idea that contrast and orientation content of the stimuli are driving the responses in early visual cortex and that coding of global properties of shape are unlikely to be found in the neural responses within these regions (Wilkinson et al., 2000).

### Stimulus properties that drive responses in object selective cortex

By contrast, tuning to low radial frequencies, whether they occurred in the RF or the RF-ripple stimuli, while rarer overall, were found consistently in object selective LO (Grill-Spector et al., 2001; Kourtzi & Kanwisher, 2000). The global properties of shape to which humans are exquisitely sensitive (Wilkinson et al., 1998) are therefore a feature of neural response in LO. LO exhibits visual field biases (Silson et al., 2016), but is not routinely identified with retinotopic mapping techniques. Indeed, in the present study, and in line with many others, LO was identified on the basis of a preference to objects over their scramble counterparts (Grill-Spector et al., 2001; Kourtzi & Kanwisher, 2000). The tunings to global shape properties are therefore consistent with the more abstract representations that have been associated with this region of the brain in previous research (Drucker & Aguirre, 2009; Haushofer et al., 2008; Op de Beeck, Haushofer, et al., 2008). It is interesting however that the two retinotopically identified regions LO-1 and LO-2 (Larsson & Heeger, 2006) also exhibit tunings to low radial frequency, which is consistent with our previous work that has shown a transition from early local processing of shape profile to processing of the complexity of shape at the same stage of the visual hierarchy (Vernon et al., 2016). The selectivity to global shape properties, which were largely maintained across our two stimuli sets, was also underscored by the multivariate patterns of response that were significantly correlated across the stimulus sets only in LO-1, LO-2 and LO.

### Is there a role for V4 in global shape processing?

V4 has been the focus of a body of research into properties that contribute to the processing of shape (see (Pasupathy et al., 2020) for a review). Work on non-human primates highlights the processing of curvature and more complex forms (Gallant et al., 1993, 1996). Recent work has led to the compelling idea that V4 is ‘shape emergent’ (Hu et al., 2020). However, studies also highlight that the curvature selective domains in V4 intermingle with domains selective to orientation and chromatic stimulus properties (Hu et al., 2020; Roe et al., 2012; Tang et al., 2020). With that backdrop it is clear that the stimuli we presented in the RF study had orientation variations that could easily have driven orientation selective responses in V4 and masked selectivity to curvature and/or global shape. Consistent with this, the responses to low radial frequencies emerged in V4 when much of the variation in orientation content of the stimuli was removed in the RF-ripple stimuli. The evidence we have gathered here therefore is consistent with V4 being a shape-emergent (Hu et al., 2020) region of the brain albeit one that processes other stimulus properties as well. There is also good evidence in human that V4 plays an important role in shape processing (Dumoulin & Hess, 2007; Wilkinson et al., 2000). The stimuli used by Dumoulin and Hess are well-suited to disentangle the contributions of orientation of stimulus elements and their arrangement into contours and this could have contributed to them showing curvature processing in V4. They also reported curvature processing in a region that likely coincided with LO-1 and LO-2.

### Areas of the brain that represents global shape defined by radial frequency

Previous work that was similar to ours has shown that radial frequency is not represented in the regions of the ventral visual pathway that we have examined here (Salmela et al., 2016). Our study differed in ways that control for variations in low level properties of radial frequency stimuli that were present in the previous study (Salmela et al., 2016) and as a result likely explains why our investigation found processing and representations of radial frequency in the regions that they did not. Here, we show that radial frequency is represented in LO on the basis of differences in radial frequency that approximate to how they are perceived – log ratios predicted the pattern of response better than differences in radial frequency. This points to LO being a region that not only processes the global curvature found in radial frequency patterns, but also represents that information in a way that has a plausible relationship with behaviour, as also demonstrated using different shape stimuli (Drucker & Aguirre, 2009; Haushofer et al., 2008; Op de Beeck, Torfs, et al., 2008). LO-1 and LO-2 also showed representations that were related well to log ratios of the radial frequency, although not as strongly as in LO. This indicates that representations of global shape emerge earlier than LO in the hierarchy of the ventral pathway, consistent with our previous work (Vernon et al., 2016).

## Conclusions

Our univariate results show clearly that two different response profiles were present in visual cortex. One showed a preference to high radial frequency patterns, which we believe are driven by neural responses to local features, while the second showed preference to low radial frequencies consistent with more global shape processing. The extent to which these response profiles were found in different regions of visual cortex varied markedly. In fact, tuning to low radial frequencies was localised to area LO and this was unaffected by the addition of high radial frequency contour modulations in the control rippled stimuli, suggesting a critical role in global shape processing. The multivariate analysis of the data also pointed to there being a global shape representation in lateral occipital regions that related well to a proxy of the perceptual differences between the stimuli, underscoring the role of lateral occipital regions in global shape processing.

## Acknowledgements

We acknowledge with thanks funding from the BBSRC (BB/P007252).

